# Stomatal conductance and net photosynthesis depend on nitrogen supply in drought-limited boreal forest

**DOI:** 10.64898/2026.11.23.768591

**Authors:** Choimaa Dulamsuren, Dugersuren Amarjargal, Dashtogtokh Bazar, Tumendelger Uitumen, Tungalag Munkhzul, Davaadorj Saindovdon, Batbold Byambatsogt, Erdenechuluun Naranbayar, Ganbaatar Batsaikhan, Markus Hauck

## Abstract

Transition from cold limitation to drought limitation due to climate change is widespread in boreal forests. We explored in the scope of the Dry Boreal Forest Long-Term Nitrogen Fertilization Experiment (DryBoreal-LTNE) in southern boreal forests of Mongolia how N supply interacts with tree water and C relations in forests of *Larix sibirica*, which are naturally drought-limited. Using branches of mature *L. sibirica* subjected to N fertilization of 6.25–25 kg N ha^−1^ a^−1^, we conducted CO_2_/H_2_O gas exchange measurements to examine how N availability interferes with the climatic control of stomatal conductance (*g*_s_) in *L. sibirica*. We tested the hypotheses that *g*_s_ increases with increasing N availability and that this increase can be translated into increased net photosynthesis (*A*) in *L. sibirica* even under the dry continental climate of Mongolia. We could confirm the first hypothesis and found that *A* decreased at low fertilization (6.25 kg N ha^−1^ a^−1^) probably caused by increases in respiration due to metabolic costs of intensified N uptake. *A* attained the highest values at 12.5 kg N ha^−1^ a^−1^, but declined at the highest fertilization level of 25 kg N ha^−1^ a^−1^. Our results demonstrate that research on the drought tolerance of boreal forests under climate change should not only focus on changes in temperature, vapor pressure deficit, and precipitation, but should take N availability into account, especially as N availability is thought to decrease in remote boreal forests due to elevated atmospheric CO_2_ concentrations and can be locally influenced by land use.

## 1. Introduction

Two paradigms of N metabolism in terrestrial ecosystems have been challenged in recent years: First, Morford et al. (2011) refuted the fundamental rule that additional reactive N (N_r_) would enter terrestrial ecosystems only from the atmosphere, when they demonstrated the uptake of rock-borne N by conifers of several species in temperate forests of northern California. Second, it has been questioned whether there is in fact a global eutrophication of ecosystems due to anthropogenic N deposition, which is widely proposed in the literature (Matson et al. 2002; Galloway et al. 2008; Rockström et al. 2024) and was thought to even concern extremely remote areas located far from N_r_ pollution sources (Holtgrieve et al. 2011). Not surprisingly, given the far-reaching implications and the complexity of the matter, challenging the second paradigm, first by case studies (McLauchlan et al. 2010, 2017) and then by more comprehensive contributions (Craine et al. 2018; Mason et al. 2022a) has initiated a not yet fully resolved debate (Craine et al. 2019; Hiltbrunner et al. 2019; Mason et al. 2022b; Olff et al. 2022).

A recent study of >1600 stemwood cores of the boreal conifers *Picea abies* and *Pinus sylvestris* from regions of different levels of N deposition in boreal and temperate Sweden provided strong support for the idea that terrestrial ecosystems outside regional hotspots of high N pollution are not only not subjected to significant eutrophication, but have even undergone a decade-long to centennial impoverishment of plant-available N (Bassett et al. 2026). This depletion in N_r_ that had already been postulated by Craine et al. (2018) and described as oligotrophication was deduced from decreasing δ^15^N signatures in tree rings since the 1950s (Bassett et al. 2026). Using mixed-effects modeling, Bassett et al. (2026) substantiated that the decrease in δ^15^N was most closely related to the atmospheric CO_2_ concentration, but was much less affected by N deposition and temperature. The significance of CO_2_ is in line with the progressive N limitation hypothesis, which postulates that N is increasingly captured in biomass under elevated atmospheric CO_2_ (eCO_2_) concentrations and thus the concentrations of available N decrease (Z. Feng et al. 2015; Liang et al. 2016), though there are also other explanations (Reich & Hobbie 2013; Craine et al. 2018; Peñuelas et al. 2020). Along with previous studies, where δ^15^N signatures were found to decrease in plant samples and tree rings since the early twentieth century (McLauchlan et al. 2010, 2017; Craine et al. 2018; Mason et al. 2022a) and where δ^15^N was shown to be positively correlated with foliar N concentrations (McLauchlan et al. 2010; Craine et al. 2018), the results of Bassett et al. (2026) suggest that oligotrophication could be widespread in the boreal forest biome.

The study of N limitation in boreal forests has a long history, as boreal forests limited by low temperatures are co-limited by low N availability, which in turn is the result of the combination of cold climate with a short growing season (Van Cleve et al. 1991; Lupi et al. 2013). Therefore, net N mineralization, the uptake of peptides and amino acids, and the effect of N fertilization have been investigated in temperature-limited boreal forests (Högberg et al. 2017). Given this situation, climate change promised improved climatic and nutritional conditions and increased productivity in most of the boreal forest biome (Huang et al. 2010; Wang et al. 2023), like in the Arctic tundra (Myers-Smith et al. 2020). However, there are also drought-limited boreal forests, which occur naturally under continental climate at the southern fringe of the biome under continental climate (Dulamsuren et al. 2023a; Wang et al. 2023), but are recently spreading over large regions due to climate change (Buermann et al. 2014; Babst et al. 2019; Mirabel et al. 2023; Sang et al. 2026). Even annual radial growth in *Larix cajanderi* trees in the earth’s coldest forests in Yakutia, where temperature minima in winter can fall far below –60°C, is today limited by drought and not by low temperatures during summer (Tei et al. 2014).

Forests in the southern boreal forest region of southern Siberia and northern Central Asia, where the boreal forest biome meets the temperate steppe grasslands of Central Asia are naturally dominantly drought-limited due to the persistent influence of the Siberian Anticyclone with its center over Lake Baikal from October to March, which shields the region from the moist westerlies for half of the year (Sahsamanoglou et al. 1991). Therefore, these forests are ideal model systems for the study of boreal forests under progressive climate change potentially representing condition that could become pervasive in the boreal forest biome. Already today boreal forests of southern Siberia and northern Central Asia are exposed to frequent and intense drought stress (Dulamsuren et al. 2019; Dulamsuren 2026), to occasional heat stress (Dulamsuren et al. 2026) and subjected to reduced growth and increased tree mortality (Dulamsuren et al. 2010, 2013; Liu et al. 2013; Kharuk et al. 2016).

Dulamsuren & Hauck (2021) used forests of *Larix sibirica* in this ecoregion as model systems to examine whether co-limitation by nitrogen, which is well-known from temperature-limited boreal forests (Högberg et al. 2017), remains effective under drought limitation. Selecting forest sites of a remote region of Mongolia with an estimated atmospheric N deposition of 1–2 kg N ha^−1^ (Rubin et al. 2023), a gradient in N supply from livestock dung was selected according to differences in densities of free-roaming livestock (Hauck & Lkhagvadorj 2013) in forests of varying stand sizes (Khansaritoreh et al. 2017). A closer relationship of foliar δ^15^N than of δ^13^C signatures to the mean sensitivity of annual stem increment and to missing ring frequency in tree-ring chronologies suggested a strong influence of N limitation on the growth of *L. sibirica* under drought (Dulamsuren & Hauck 2021). Like in the time series of McLauchlan et al. (2010) and Craine et al. (2018), decreases in δ^15^N in *L. sibirica* (analyzed in needles) were reflected by decreasing N content in the comparison of sites with different N supply. The significance of N co-limitation also for strongly drought-limited boreal forests under the ultracontinental climate of Mongolia is consistent with the known decrease of net N mineralization and organic matter decomposition under drought (Peñuelas et al. 2020) and suggests that drought and N limitation can be expected to concur in the boreal forest biome in general, where low temperature limitation is not or no longer dominant.

There are different mechanisms how the N supply affects water and C relations in trees. In addition to biomass increases due to the fertilization effect, high N availability causes shifts towards lowered root/shoot ratios (Ågren & Franklin 2003; H. Feng et al. 2023; F. Feng et al. 2026a), influences fine-root morphology and mycorrhization (Dietrich et al. 2025), and interferes with the drought regulation of stomatal control (Guo et al. 2003; Hauck & Dietrich 2025). Greater investments in aboveground biomass at the cost of a less extensively developed root system follow the optimal partitioning theory (Kobe et al. 2010), as less resources are needed to acquire N from the soil, but increase the risk that the root system is not capable of absorbing enough water during drought (Dziedek et al. 2016). Because *L. sibirica* has generally a small fine-root biomass, which has been shown to be even further reduced under drought (Chenlemuge et al. 2013), a major reduction of the already low fine-root biomass due to high N availability seems rather unlikely in this species, but has not yet been investigated.

High N supply can decouple the control of stomatal conductance (*g*_s_) from soil moisture and the atmospheric vapor pressure deficit (VPD), because NO_3_^−^ stimulates stomatal opening if accumulated in the stomatal guard cells (Guo et al. 2003). NO_3_^−^ uptake in the guard cells is regulated by specific transporters (Tsay et al. 2007) and not influenced by abscisic acid (ABA), which mediates the drought control of *g*_s_ in response to soil moisture (Merilo et al. 2015; Hsu et al. 2021). High foliar NO_3_^−^ concentrations are thus capable to counteract drought-induced stomatal closure signaled by ABA. This way, NO ^−^ forces plants to a riskier drought response, which is untypical for a Pinaceae species, like *L. sibirica* (Dulamsuren et al. 2009; Dulamsuren 2026) and most other boreal conifers (Brodribb et al. 2014). At moderate drought, the NO_3_^−^-mediated less stringent stomatal control can be beneficial, as increased *g*_s_ allows for increases in CO_2_ assimilation. The greater interannual growth variability (indicated by higher mean sensitivity) in the tree-ring data of *L. sibirica* from N-rich sites found by Dulamsuren & Hauck (2021) was attributed to faster growth during moderate drought. If a drought intensifies, the loss of plant water due to the extended stomatal opening, however, might become critical resulting in the increased missing ring frequency observed in *L. sibirica* in N-enriched forests (Dulamsuren & Hauck 2021) and finally increase tree mortality.

While combined data of stable isotopes and biomass increment are supportive of the existence of a causal relationship of N supply on tree growth via an influence on stomatal control in *L. sibirica* (Dulamsuren & Hauck 2021), no data were available that directly demonstrated an effect of N availability on *g*_s_ in *L. sibirica* or any other tree species from a drought-limited boreal forest. Therefore, and for the study of other causal effects of N in drought-limited boreal forests, we established the Dry Boreal Forest Long-Term Nitrogen Fertilization Experiment (DryBoreal-LTNE) in *L. sibirica* forests of Mongolia.

To directly demonstrate the relationship of N supply with *g*_s_ in drought-limited boreal forests and on parameters of the CO_2_/H_2_O exchange, which depend on *g*_s_ – i.e. transpiration (*E*), net photosynthesis (*A*), water-use efficiency (WUE) – we analyzed the effect of N fertilization on the CO_2_/H_2_O exchange in mature *L. sibirica* trees. Because the CO_2_/H_2_O exchange is highly sensitive to temperature (Busch et al. 2024) and the climate in the boreal forest biome is rapidly warming (Gauthier et al. 2015), we also varied temperature in a range of 22–40°C, which is realistic for ambient canopy temperatures in *L. sibirica* forests (Dulamsuren et al. 2026). We aim at testing the hypotheses (1) that increasing N availability increased both *g*_s_ and *E* and that (2) the assumed increase in *g*_s_ paid off in the form of increased *A* despite the increase in dark respiration (*R*_d_) that had to be expected during increased N uptake. The second hypothesis was based on the published fact that spatially varying N availability in different *L. sibirica* stands resulted in differences in the interannual variability of radial stem increment (Dulamsuren & Hauck 2021). Assuming that the improved N supply allows for increased investments in the photosynthetic apparatus (e.g. increased synthesis of Rubisco and chlorophylls), we also tested the hypothesis (3) that N fertilization resulted in increases in the photosynthetic capacity (*A*_max_) and water-use efficiency (WUE).

## 2. Materials and methods

### 2.1. Study area

The study was carried out in drought-limited southern boreal forests of *Larix sibirica* Ledeb. in the Central Asian/southern Siberian boreal forest region. The studied forests are located in the Khangai Mountains of northern-central Mongolia near the town of Jargalant (48°44’ N, 100°46’ E, 1600 m a.s.l.), Arkhangai district. Forests in the Khangai Mountains form the southernmost distribution limit of boreal forest in Mongolia and largely limited to the cooler and moister north-facing slopes in this region. All stands were natural mature forests without systematic forest management, but sporadic utilization for selective logging of dead trees.

### 2.2. Dry Boreal Forest Long-Term Nitrogen Fertilization Experiment (DryBoreal-LTNE)

The Dry Boreal Forest Long-Term Nitrogen Fertilization Experiment (DryBoreal-LTNE) is designed as a long-term N fertilization experiment, which has been set up by the Chair of Applied Vegetation Ecology (Faculty of Environment and Natural Resources, University of Freiburg, Germany) in 2024 (Table 1). It consists of an unfertilized control (treatment N0; 0 kg N ha^−1^ a^−1^) and three N fertilization treatments (N1 to N3 with 6.25, 12.5, and 25 kg N ha^−1^ a^−1^). The experiment is replicated five times on the plot level, with 5 clusters of N0 to N3 plots distributed over north-facing mountain slopes of a rectangular research area of 15 km in latitude × 5 km in longitude at a mean distance between study sites of 8.7±1.9 km and at an elevation of 1725–1840 m a.s.l. Each of the 20 plots has a size of 20 m × 20 m. The clusters of plots are termed “site” in the analyses.

**Table 1.** Design of Dry Boreal Forest Long-Term Nitrogen Fertilization Experiment (DryBoreal-LTNE).

| Category | Specification |
|---|---|
| **Treatments** |  |
| N0 | 0 kg N ha<sup>-1</sup> a<sup>-1</sup> |
| N1 | 6.25 kg N ha<sup>-1</sup> a<sup>-1</sup> |
| N2 | 12.5 kg N ha<sup>-1</sup> a<sup>-1</sup> |
| N3 | 25 kg N ha<sup>-1</sup> a<sup>-1</sup> |
| **Application of fertilizer** | Urea dissolved in 120 l water per plot and treatment (N0 only water)<br>Two treatments per year with 0, 3.125, 6.25, 12.5 kg N ha<sup>-1</sup> June and August |
| **Start of experiment** | June 2024 |
| **Replication** | 5 clusters of N0–N3 plots of 20 m × 20 m |

The fertilizer is applied as urea (CH_4_N_2_O). Half of the annual dose is applied in June and the other half in August. The growing season lasts from late May to September. The urea is dissolved in 120 l of water per plot. On the control plots (N0), 120 l without fertilizer are applied to avoid that data are biased by an irrigation effect. We conducted the gas exchange measurements in the third year of N fertilization.

### 2.3. Climate

Climate of the study area was characterized based Copernicus ERA5 re-analysis data (https://cds.climate.copernicus.eu) for January 1940 to August 2026 (Fig. 1). Monthly mean air temperature at 2 m and monthly sums of precipitation were averaged for the grid field of 48.5 – 49.0 °N and 100.5 – 101.0 °E. The download of ERA5 data in grib format was done with the R package ‘ecmfr’ 2.0.3 and data processing and conversion of grib files was done with the R package ‘terra’ 1.9-11. Precipitation sums were corrected for overestimation inherent in the ERA5 data for summer precipitation in continental areas (Lavers et al. 2022; Soci et al. 2025) by linear regression with CRU TS 4.04 data edited by the Climate Research Unit of the University of East Anglia, Norwich and the Met Office in Exeter, UK (https://climexp.knmi.nl) for 1940–2023 using the function *y* = 0.679*x*-0.6466.

**Fig. 1.**
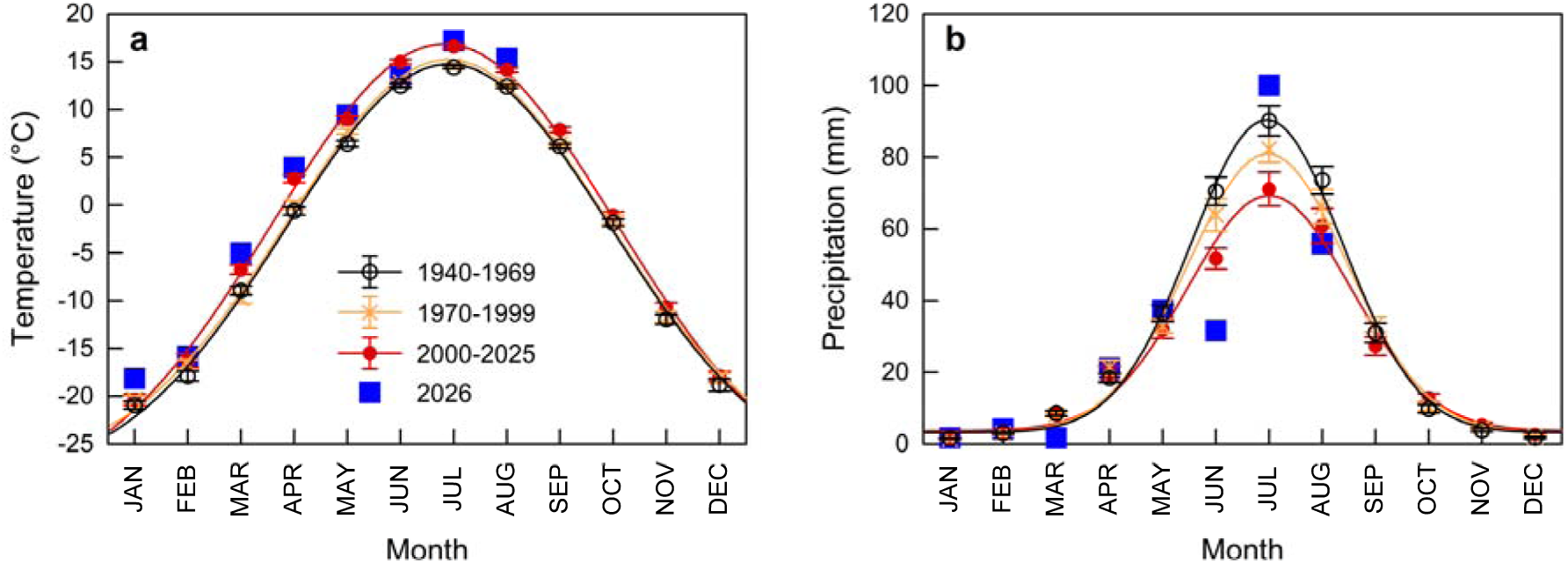
Temporal trends of monthly air temperature and precipitation (1940–2026) for the study area from ERA5 data: (a) Monthly mean air temperature at 2 m (±standard errors) and (b) monthly sums of precipitation averaged for the grid field of 48.5 – 49.0 °N and 100.5 – 101.0 °E based on Copernicus ERA5 data (https://cds.climate.copernicus.eu) for January 1940 to August 2026. Precipitation sums were corrected for overestimation inherent in the ERA5 data for summer precipitation in continental areas (Lavers et al., 2022; Soci et al., 2025) by linear regression with CRU TS 4.04 data edited by the Climate Research Unit of the University of East Anglia, Norwich and the Met Office in Exeter, UK (https://climexp.knmi.nl) for 1940–2023 using the function *y* = 0.679*x*-0.6466.

Stand microclimate below the canopy (Fig. 2) was recorded every 1 h from every research plot (*N*=20) with three randomly distributed with HOBO U23 ProV2 temperature/relative humidity data loggers (Onset Computer Cooperation, Bourne, Massachusetts, USA). These loggers were mounted at a height of 2 m at the north side of *L. sibirica* trees. Records were available from August 2024 to August 2026. Since our CO_2_/H_2_O gas exchange measurements were done in June and August 2026, we have limited data presentation microclimate to the summer months (June to August) within the available dataset. Maximum canopy temperatures have been shown to exceed the below-canopy temperatures by 8.4±2.5 K based on drone flights conducted for thermal imaging in the stands of DryBoreal-LTNE in June, July, and August 2025 (Dulamsuren et al. 2026). VPD (in kPa) was calculated from air temperature (*T*) and relative humidity (RH) using the Magnus-Tetens equation VPD = (0.61078 kPa × exp((17.27 *T*)/(*T* + 237.3))) × (1 - (RH/100)) for vapor pressures over fluid water (Anyadike 1984).

**Fig. 2.**
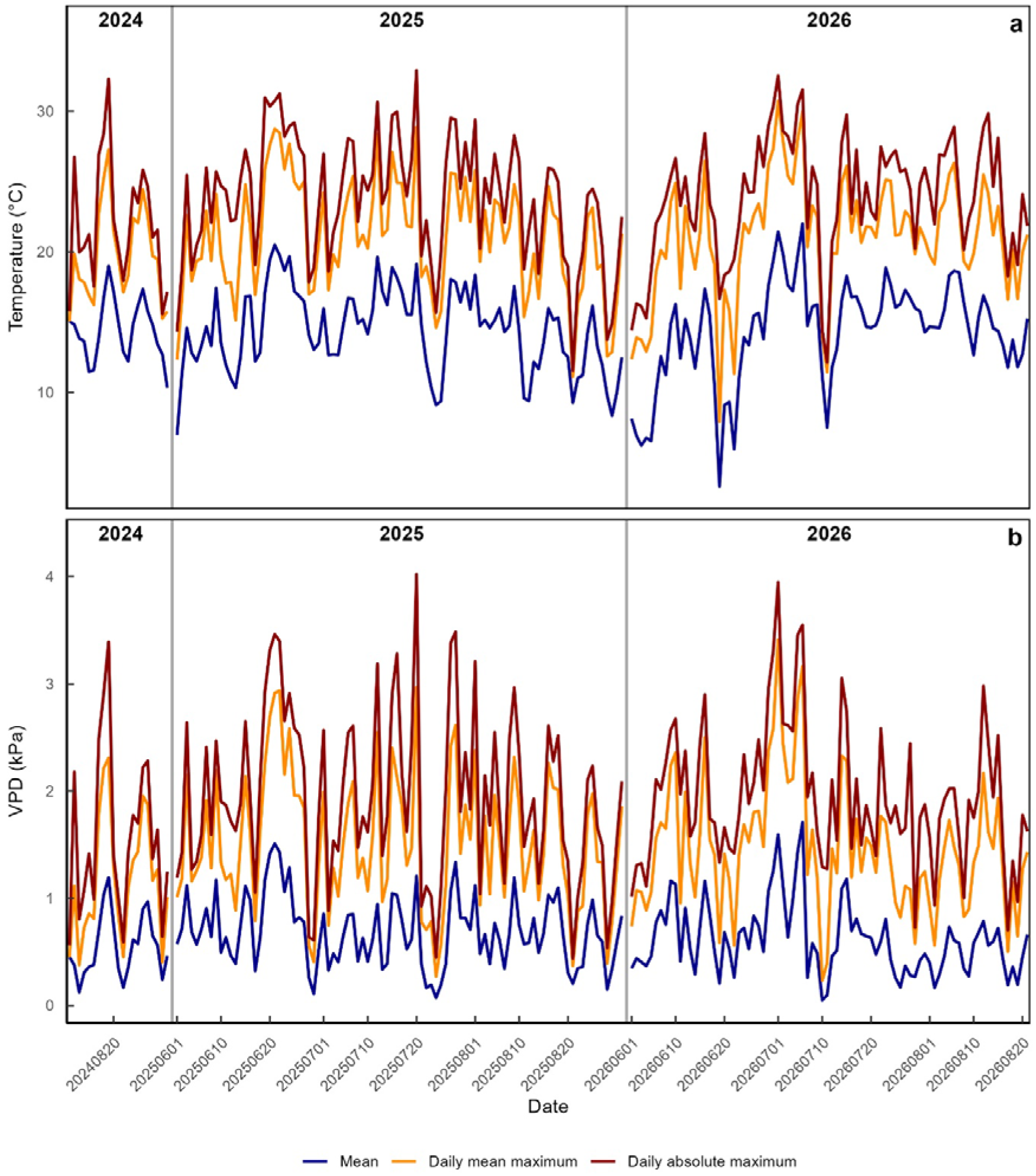
Stand microclimate measured below the canopy at 2 m height from August 2024 to August 2026. Only summer climate from June to August is displayed: (a) air temperature, (b) VPD. Values based on 3 loggers on 20 plots including 5 sites each with 4 fertilization treatments (N0–N3).

Mean annual temperature from 1940–2025 amounted to −1.7±0.1°C (Fig. 1a) and has increased at a rate of 0.31 K decade^−1^ since 1940 and 0.34 K decade^−1^ since 1970. Mean July and January temperatures from 1940–2025 amounted to 15.1±0.2°C and −20.6±0.3°C. Mean annual precipitation from 1940–2025 was 327±6 mm a^−1^ (Fig. 1b). The precipitation maximum in summer has been markedly reduced (Fig. 1b). Based on the linear regression line, mean annual precipitation has declined from 361 mm a^−1^ to 293 mm a^−1^ by 68 mm or at a rate of 8 mm decade^−1^. Precipitation in June 2026 was only 51% of the long-term mean of 1940–2025 and 61% of the mean of 2000–2025. July precipitation of 2026 was above the long-term mean (122% of mean 1940–2025; 141% of mean 2000–2025). The August of 2026 was slightly drier than the average (83% of mean 1940–2025; 92% of mean 2000–2025).

Mean below-canopy temperatures (Fig. 2) recorded in the summer months (June to August) from August 2024 to August 2026 in the studied *L. sibirica* stands was 14.4±0.2°C. Maximum temperatures amounted to 21.0±0.3°C if calculated as the daily mean maximum of the 60 loggers that were distributed over the 20 sample plots, and 23.7±0.3°C if the daily absolute maximum recorded in at least one of the loggers was selected. The absolute highest below-canopy air temperature values were 30.7°C (daily mean maximum across all loggers) and 32.9°C (absolute maximum). Based on Dulamsuren et al. (2026), this corresponded to peaks in the canopy temperature of 39.1 and 41.3°C.

### 2.4. CO_2_/H_2_O gas exchange measurements

Sun-exposed branches of mature *L. sibirica* trees were collected with a Notch Big Shot slingshot (Notch Equipment, Greensboro, North Carolina, USA) from three replicate trees per research plot. Branches were allowed to saturate with water for at least 1 day by putting them into a water bucket under dim light. Air humidity was not controlled during this pretreatment. Storage between sampling and measurement was for 2.8±0.1 days. The potential influence of the number of storage days was controlled for by including it as a random factor in all our regression models. The residual variance that was attributable to the number of storage days and the generally moderate differences between conditional and marginal *R*^2^ demonstrated a generally low effect of storage in results, which was nevertheless consistently included in modeling by using mixed-effects models throughout. The effect on *E* was moderate, but higher than on the other variables.

Photosynthetic photon flux density (PPFD) was varied between 0, 20, 40, 60, 80, 100, 150, 300, 600, 900, 1200, and 1500 µmol m^−2^ s^−1^. We started with the highest light intensity and regulated stepwise down to dark to allow the plant to acclimate to the highest light intensity during the stabilization phase of leaf gas exchange in the measuring chamber, before the light-response curve was recorded. Stabilization was allowed for a minimum of 15 min, but this period was extended for many samples based on careful visual control of the stability of *A* and *g*_s_ for each individual sample, before the measurement was started.

Four light-response curves at an air temperature of 22, 30, 35, and 40°C were recorded consecutively in the same sample. To avoid that samples were affected from photoinhibition from the previous measurement, we decided to limit the maximum light intensity to 1500 µmol m^−2^ s^−1^. This is below the highest values of solar irradiation that are to be expected under bright sunlight in the sun crown of *L. sibirica* forests under the highly continental climate of Mongolia with 250–260 sunny day per year (Yembuu 2021). However, 1500 µmol m^−2^ s^−1^ are already sufficient to determine *A*_max_ from the fitted curve. Our solution is thus a compromise, as we otherwise would have been urged to take new subsamples for the individual light-response curves at each temperature otherwise, which might have cause additional variance in the dataset. Moreover, we considered our design with a ramped temperature increase from light curve to light curve more realistic, as high canopy temperatures of 40°C can be expected to be gradually built up during a heat wave. Relative humidity was set to 30% taking into account Mongolia’s dry and highly continental climate combined with the fact that we examined branches that were sun-exposed in the field. CO_2_ concentration was set to 400 ppm.

Needles were detached from the shoots after the experiment, air-dried and then dried at 105°C for at least 24 h and weighed. All specifications in the paper are per g dry weight. The weight of 50 fresh randomly selected needles was 62.692±1.877 mg (*N*=5), which enables the approximate conversion to leaf area-based data if desired for comparisons.

### 2.5. Statistical analyses

All statistical analyses were computed in R 4.4.1. Arithmetic means ± standard errors are given throughout the paper. *P* values of significant effects (*P*≤0.05) in the tables are printed in bold.

#### 2.5.1. Light-response curve fitting

For fitting the light-response curves of *A* vs. PPFD, we did not use the often-recommended non-rectangular hyperbola (He et al. 2024), but the more robust rectangular hyperbola (Gomes et al. 2006), because we limited the maximum of the PPFD gradient to 1500 µmol m^−2^ s^−1^. Since we collected the branches used for the measurements from the true sun-exposed sun crown and the continental climate of Mongolia often generates high solar irradiance, 1500 µmol m^−2^ s^−1^ was only at the onset of light saturation. The additional curvature parameter (θ) in non-rectangular models (Thornley 1998), therefore, led to implausible results in some curves with very sharp increases in *A* due to overfitting. To overcome this problem and because we selected our experimental design with maximum PPFD of 1500 µmol m^−2^ s^−1^ intentionally, we decided for curve fitting with the rectangular hyperbola. The model selection for light-response curve fitting can affect the absolute values of *A*_max_, LCP, and *R*_d_, but not the values of *A* and WUE used in our regression models.

Curve fitting with the rectangular hyperbola was done using the equation *A* = (*a* PPFD)/(*b* + PPFD) + *c* and was calculated with the nonlinear least squares (nls) function of R. In this equation, *a* stands for *A*_max_, *c* for *R*_d_, and *b* represents the PPFD value where *A* is 50% of *A*_max_ (Lobo et al. 2013).

#### 2.5.2. Regression modeling

Statistical analyses were based on the calculation of linear mixed-effects models using the R packages ‘lme4’ 1.1-35.5 and ‘lmerTest’ 3.2-1 as well as the ‘performance’ 0.17.0 and ‘DHARMa’ 0.4.6 for quality checks. The reason for choosing mixed-effects models was to control for potential influences of the site and of sample storage time before the measurement, which were both included as random factors. These random factors had a low influence in many cases, as was indicated by near-identical conditional and marginal *R*^2^ values, but we kept the mixed-effects models consistently and did not switch to simple regression models even in these cases to document the effects or non-effects quantitatively. The independent parameters in the model did not show any multicollinearity if analyzed without interactions. Interactions between parameters were only examined if the variance inflation factor was ≤4.

## 3. Results

### 3.1. Response to nitrogen

Linear mixed-effects modeling revealed positive effects of increasing N fertilization on both *g*_s_ and *E* (Table 2). The effect of N was small compared to other independent parameters, but significant. N fertilization also increased *A* and this effect was stronger than that on *g*_s_ and *E* (Table 2). A comparison of the light saturation curves of *A* across the different N fertilization levels showed that the increase of *A* with N availability was not steady (Fig. 3). Rather, *A* decreased first at the lowest fertilization level N1 of 6.25 kg N ha^−1^ a^−1^, but increased strongly to higher values than the control N0 (0 kg N ha^−1^ a^−1^) in the N2 treatment (12.5 kg N ha^−1^ a^−1^). At the highest level of fertilization N3 (25 kg N ha^−1^ a^−1^), *A* decreased to values below those in N2 and were only slightly higher than or even identical with the *A* values in N0. The increase of *A* in N2 compared to N0 was significant in the linear mixed-effects model (Table 3). The response of *A* to N fertilization was blurred at high temperatures of 35–40°C (Fig. 3). This was reflected by a positive interaction of N1 × temperature, which indicated that the negative effect of low N fertilization was reduced at high temperatures (Table 3).

**Fig. 3.**
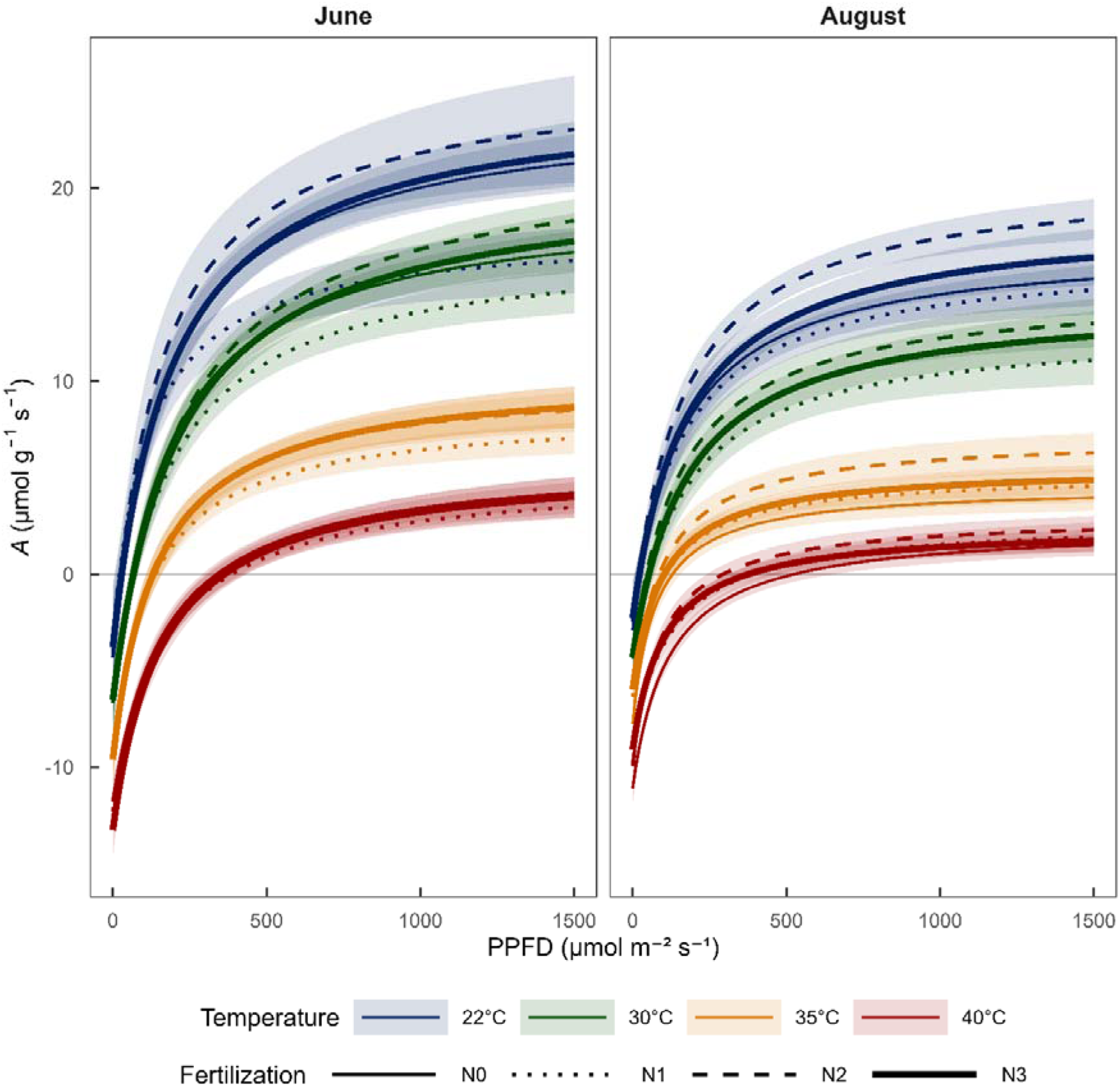
Light response of net photosynthesis (*A*) in *Larix sibirica* at different temperatures from 22–40°C in June and August in the third year of fertilization with 0 (N0), 6.25 (N1), 12.5 (N2), and 25 (N3) kg N ha^−1^ a^−1^.

**Table 2.**
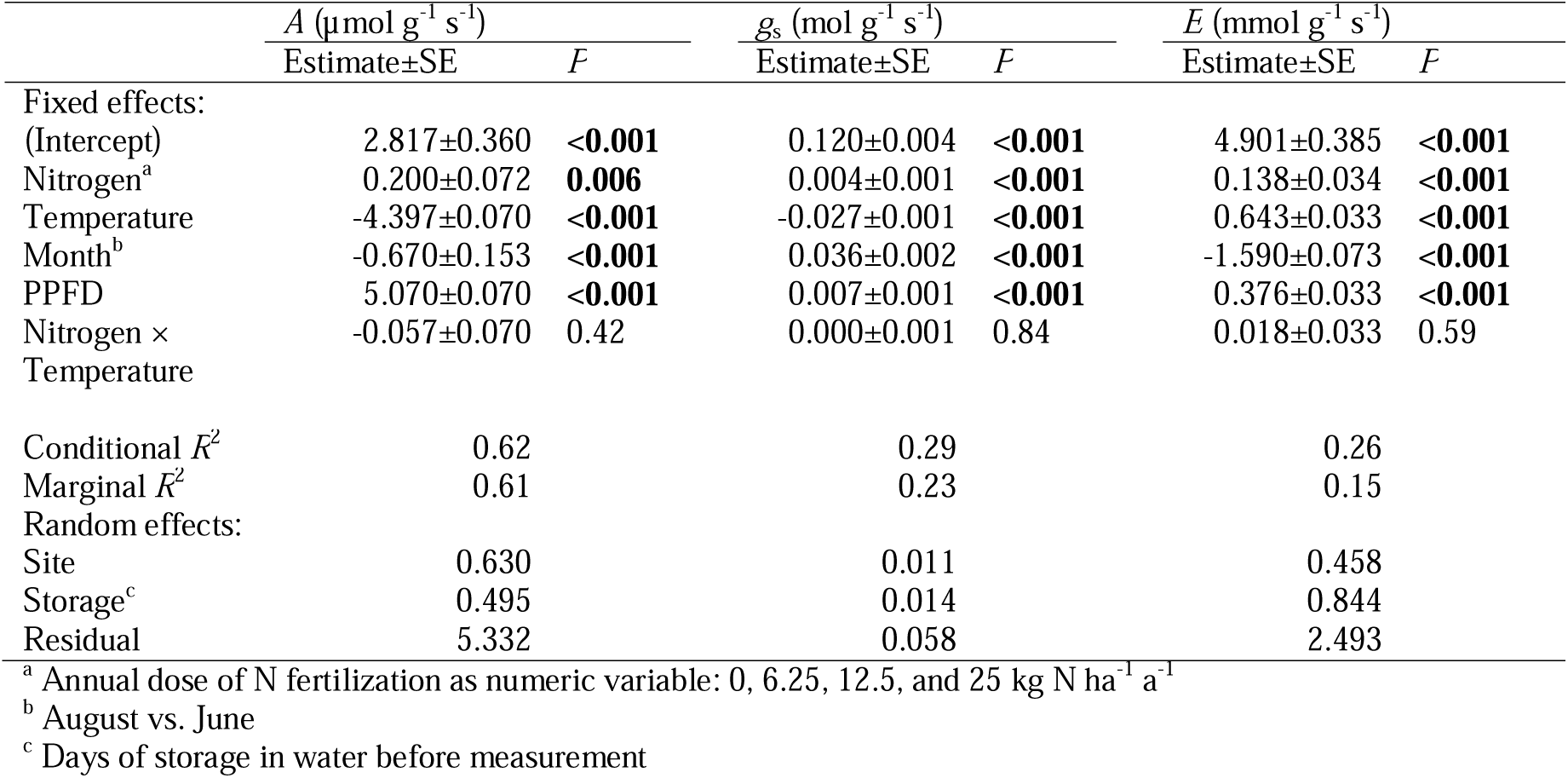
Linear mixed-effects model analyzing effects of N fertilization, temperature, and month on net photosynthesis (*A*), stomatal conductance (*g*_s_), transpiration (*E*) in *Larix sibirica*.

|  | *A* (µmol g<sup>-1</sup> s<sup>-1</sup>) |  | *g*<sub>s</sub> (mol g<sup>-1</sup> s<sup>-1</sup>) |  | *E* (mmol g<sup>-1</sup> s<sup>-1</sup>) |  |
|---|---|---|---|---|---|---|
|  | Estimate±SE | *P* | Estimate±SE | *P* | Estimate±SE | *P* |
| Fixed effects: |  |  |  |  |  |  |
| (Intercept) | 2.817±0.360 | <**0.001** | 0.120±0.004 | <**0.001** | 4.901±0.385 | <**0.001** |
| Nitrogen<sup>a</sup> | 0.200±0.072 | **0.006** | 0.004±0.001 | <**0.001** | 0.138±0.034 | <**0.001** |
| Temperature | -4.397±0.070 | <**0.001** | -0.027±0.001 | <**0.001** | 0.643±0.033 | <**0.001** |
| Month<sup>b</sup> | -0.670±0.153 | <**0.001** | 0.036±0.002 | <**0.001** | -1.590±0.073 | <**0.001** |
| PPFD | 5.070±0.070 | <**0.001** | 0.007±0.001 | <**0.001** | 0.376±0.033 | <**0.001** |
| Nitrogen × Temperature | -0.057±0.070 | 0.42 | 0.000±0.001 | 0.84 | 0.018±0.033 | 0.59 |
| Conditional *R*<sup>2</sup> | 0.62 |  | 0.29 |  | 0.26 |  |
| Marginal *R*<sup>2</sup> | 0.61 |  | 0.23 |  | 0.15 |  |
| Random effects: |  |  |  |  |  |  |
| Site | 0.630 |  | 0.011 |  | 0.458 |  |
| Storage<sup>c</sup> | 0.495 |  | 0.014 |  | 0.844 |  |
| Residual | 5.332 |  | 0.058 |  | 2.493 |  |

<sup>a</sup> Annual dose of N fertilization as numeric variable: 0, 6.25, 12.5, and 25 kg N ha<sup>-1</sup> a<sup>-1</sup>
<sup>b</sup> August vs. June
<sup>c</sup> Days of storage in water before measurement

**Table 3.**
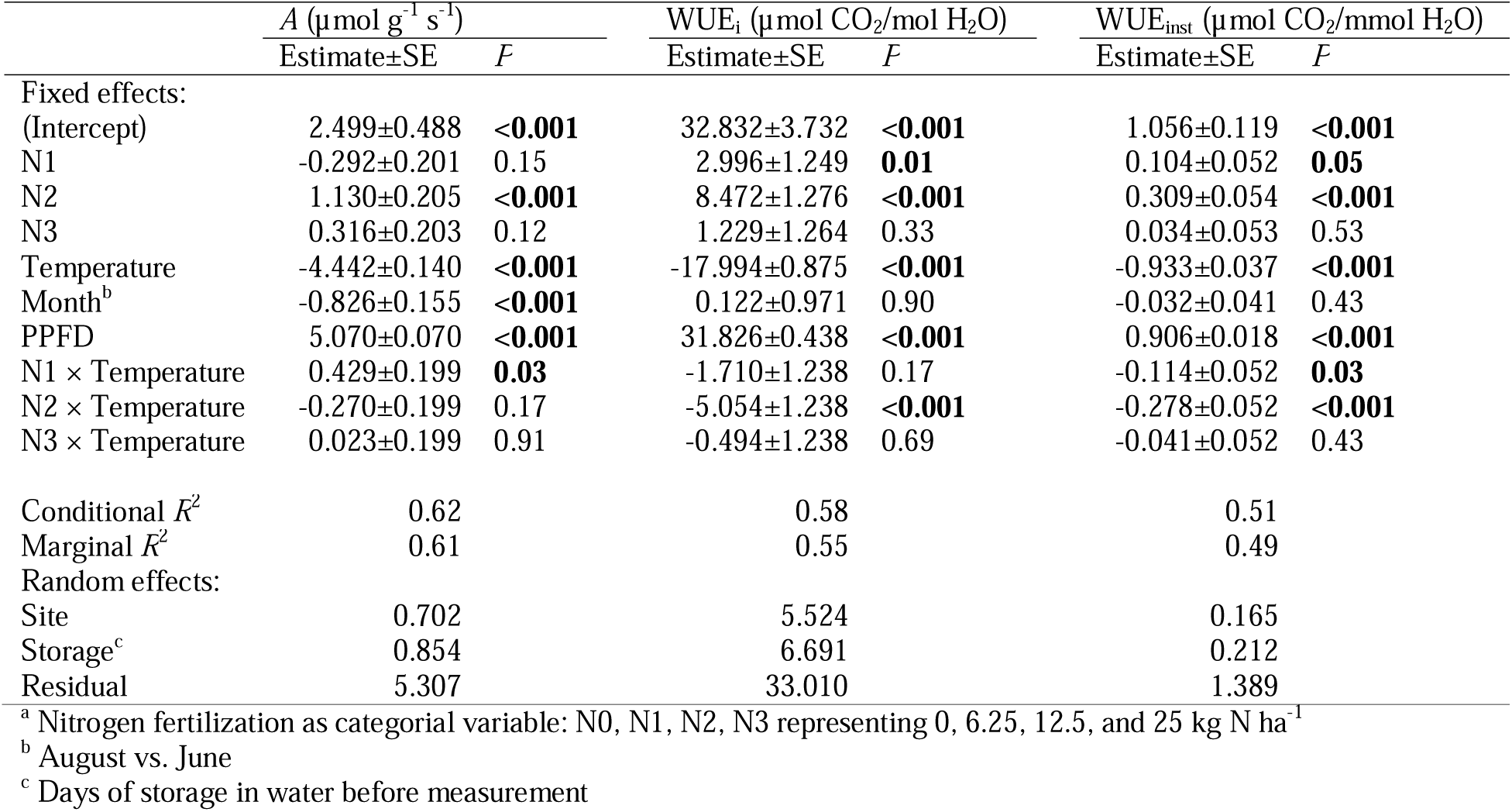
Linear mixed-effects model analyzing effects of the N fertilization treatment (variants N0–N3)^a^, temperature, and month on net photosynthesis (*A*) and on on intrinsic (WUE_i_) and instantaneous (WUE_inst_) water-use efficiency in *Larix sibirica*.

|  | *A* (µmol g<sup>-1</sup> s<sup>-1</sup>) |  | WUE<sub>i</sub> (µmol CO<sub>2</sub>/mol H<sub>2</sub>O) |  | WUE<sub>inst</sub> (µmol CO<sub>2</sub>/mmol H<sub>2</sub>O) |  |
|---|---|---|---|---|---|---|
|  | Estimate±SE | *P* | Estimate±SE | *P* | Estimate±SE | *P* |
| Fixed effects: |  |  |  |  |  |  |
| (Intercept) | 2.499±0.488 | <**0.001** | 32.832±3.732 | <**0.001** | 1.056±0.119 | <**0.001** |
| N1 | -0.292±0.201 | 0.15 | 2.996±1.249 | **0.01** | 0.104±0.052 | **0.05** |
| N2 | 1.130±0.205 | <**0.001** | 8.472±1.276 | <**0.001** | 0.309±0.054 | <**0.001** |
| N3 | 0.316±0.203 | 0.12 | 1.229±1.264 | 0.33 | 0.034±0.053 | 0.53 |
| Temperature | -4.442±0.140 | <**0.001** | -17.994±0.875 | <**0.001** | -0.933±0.037 | <**0.001** |
| Month<sup>b</sup> | -0.826±0.155 | <**0.001** | 0.122±0.971 | 0.90 | -0.032±0.041 | 0.43 |
| PPFD | 5.070±0.070 | <**0.001** | 31.826±0.438 | <**0.001** | 0.906±0.018 | <**0.001** |
| N1 × Temperature | 0.429±0.199 | **0.03** | -1.710±1.238 | 0.17 | -0.114±0.052 | **0.03** |
| N2 × Temperature | -0.270±0.199 | 0.17 | -5.054±1.238 | <**0.001** | -0.278±0.052 | <**0.001** |
| N3 × Temperature | 0.023±0.199 | 0.91 | -0.494±1.238 | 0.69 | -0.041±0.052 | 0.43 |
| Conditional *R*<sup>2</sup> | 0.62 |  | 0.58 |  | 0.51 |  |
| Marginal *R*<sup>2</sup> | 0.61 |  | 0.55 |  | 0.49 |  |
| Random effects: |  |  |  |  |  |  |
| Site | 0.702 |  | 5.524 |  | 0.165 |  |
| Storage<sup>c</sup> | 0.854 |  | 6.691 |  | 0.212 |  |
| Residual | 5.307 |  | 33.010 |  | 1.389 |  |

<sup>a</sup> Nitrogen fertilization as categorial variable: N0, N1, N2, N3 representing 0, 6.25, 12.5, and 25 kg N ha<sup>-1</sup>
<sup>b</sup> August vs. June
<sup>c</sup> Days of storage in water before measurement

*A*_max_ and *R*_d_ showed no steady response to increasing N supply (Table S1). It increased significantly compared to the unfertilized control (N0) in the N2 treatment, whereas N1 and N3 had no significant effect on *A*_max_ (Table 4). *R*_d_ was significantly intensified (indicated by more strongly negative values) in the N1 treatment (Table 4), though it should be noted that the model for *R*_d_ had a large residual variance and thus a low *R*^2^, indicating that the control of *R*_d_ was complex and only partly covered by N and temperature. LCP was not affected by N fertilization, neither in a model with the N fertilization treatment as a categorial variable (not shown) nor in a model with the N dose as numeric parameter (Table S1).

**Table 4.** Linear mixed-effects model analyzing effects of the N fertilization treatment (variants N0–N3)^a^, temperature, and month on relative values of the photosynthetic capacity (*A*_max_) in *Larix sibirica* where *A*_max_ and *R*_d_ at 22°C were set to 100% and *A*_max_ and *R*_d_ at the other temperatures were related to these values in percent.

|  | *A*<sub>max</sub> (µmol g<sup>-1</sup> s<sup>-1</sup>) |  | *R*<sub>d</sub> (µmol g<sup>-1</sup> s<sup>-1</sup>) |  |
|---|---|---|---|---|
|  | Estimate±SE | *P* | Estimate±SE | *P* |
| Fixed effects: |  |  |  |  |
| (Intercept) | 61.352±4.811 | <**0.001** | 297.88±148.39 | 0.06 |
| N1 | 8.552±5.692 | 0.13 | -342.24±141.99 | **0.021** |
| N2 | 12.045±5.721 | **0.04** | -23.40±143.44 | 0.87 |
| N3 | 7.505±5.742 | 0.20 | 25.99±143.88 | 0.86 |
| Temperature | -35.872±4.029 | <**0.001** | 106.13±100.92 | 0.29 |
| Month<sup>b</sup> | -14.780±4.115 | <**0.001** | -76.94±104.77 | 0.46 |
| N1 × Temperature | 5.904±5.698 | 0.30 | -209.41±142.72 | 0.14 |
| N2 × Temperature | 11.345±5.698 | **0.05** | -11.25±142.72 | 0.94 |
| N3 × Temperature | 7.467±5.699 | 0.19 | 37.62±141.41 | 0.79 |
| Conditional *R*<sup>2</sup> | 0.34 |  | 0.07 |  |
| Marginal *R*<sup>2</sup> | 0.33 |  | 0.03 |  |
| Random effects: |  |  |  |  |
| Site | 0.000 |  | 152.82 |  |
| Storage<sup>c</sup> | 3.672 |  | 155.84 |  |
| Residual | 43.932 |  | 1092.98 |  |

<sup>a</sup> Nitrogen fertilization as categorial variable: N0, N1, N2, N3 representing 0, 6.25, 12.5, and 25 kg N ha<sup>-1</sup>
<sup>b</sup> August vs. June
<sup>c</sup> Days of storage in water before measurement

Both WUE_i_ and WUE_inst_ were significantly increased by N fertilization in N1 and even more in N2 (Table 3). At the highest N fertilization level N3, WUE_i_ and WUE_inst_ were not significantly different from N0 (Table 3). The positive effects of N supply on WUE were reduced at high temperature, as became evident from significant negative interaction of N2 × temperature for WUE_i_ and WUE_inst_ and of N1 × temperature for WUE_inst_ (Table 3). If N fertilization was included as a numeric variable, there was no significant effect of the annual N dose on WUE_i_ and WUE_inst_, indicating that the influence of N fertilization on WUE was not a steady increase (Table S2).

### 3.2. Effects of high temperature

Increasing temperature had an overall negative effect on *g*_s_ and *A*, but a positive effect on *E* (Table 2; Fig. 3). The magnitude of the effects of the temperature, which was varied for each individual branch during the measurements, was higher than the effect of N fertilization in the years before the measurements. Even though *g*_s_ and *E* showed contrasting responses to temperature, both WUE_i_ (calculated from *g*_s_) and WUE_inst_ (calculated from *E*) decreased with increasing temperature (Table 3). Mixed-effects model were temperature was treated as a categorial variable demonstrated that the decrease of *A* with increasing temperature was steady, whereas *g*_s_ was steadily reduced from 22–35°C, but was slightly higher at 40°C than at 35°C (Table 5).

**Table 5.**
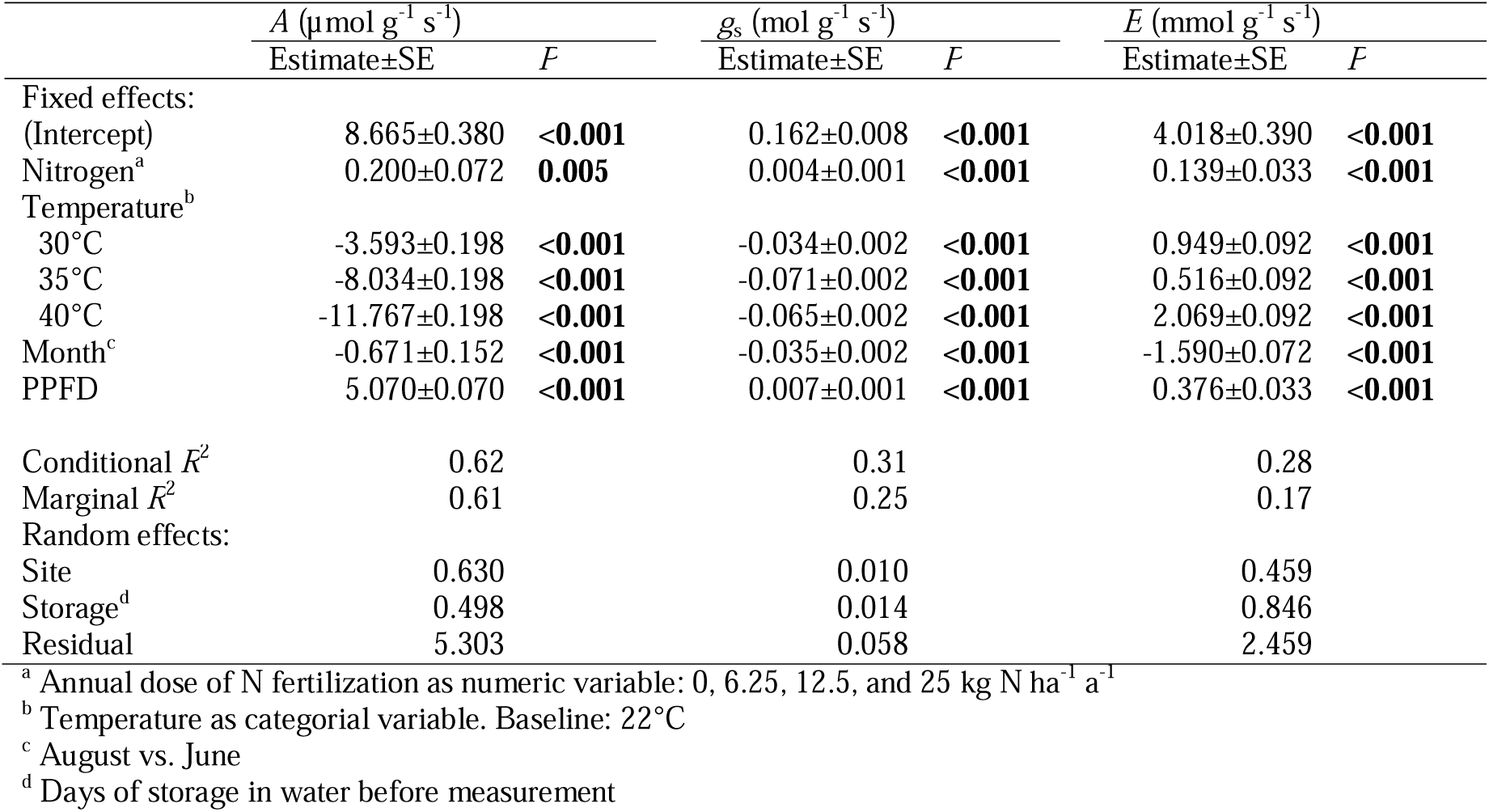
Linear mixed-effects model analyzing effects of the individual temperature variants, N fertilization, temperature, and month on net photosynthesis (*A*), stomatal conductance (*g*_s_), transpiration (*E*) in *Larix sibirica*.

|  | *A* (µmol g<sup>-1</sup> s<sup>-1</sup>) |  | *g*<sub>s</sub> (mol g<sup>-1</sup> s<sup>-1</sup>) |  | *E* (mmol g<sup>-1</sup> s<sup>-1</sup>) |  |
|  | Estimate±SE | *P* | Estimate±SE | *P* | Estimate±SE | *P* |
|---|---|---|---|---|---|---|
| Fixed effects: |  |  |  |  |  |  |
| (Intercept) | 8.665±0.380 | **<0.001** | 0.162±0.008 | **<0.001** | 4.018±0.390 | **<0.001** |
| Nitrogen<sup>a</sup> | 0.200±0.072 | **0.005** | 0.004±0.001 | **<0.001** | 0.139±0.033 | **<0.001** |
| Temperature<sup>b</sup> |  |  |  |  |  |  |
| 30°C | -3.593±0.198 | **<0.001** | -0.034±0.002 | **<0.001** | 0.949±0.092 | **<0.001** |
| 35°C | -8.034±0.198 | **<0.001** | -0.071±0.002 | **<0.001** | 0.516±0.092 | **<0.001** |
| 40°C | -11.767±0.198 | **<0.001** | -0.065±0.002 | **<0.001** | 2.069±0.092 | **<0.001** |
| Month<sup>c</sup> | -0.671±0.152 | **<0.001** | -0.035±0.002 | **<0.001** | -1.590±0.072 | **<0.001** |
| PPFD | 5.070±0.070 | **<0.001** | 0.007±0.001 | **<0.001** | 0.376±0.033 | **<0.001** |
| Conditional *R*<sup>2</sup> | 0.62 |  | 0.31 |  | 0.28 |  |
| Marginal *R*<sup>2</sup> | 0.61 |  | 0.25 |  | 0.17 |  |
| Random effects: |  |  |  |  |  |  |
| Site | 0.630 |  | 0.010 |  | 0.459 |  |
| Storage<sup>d</sup> | 0.498 |  | 0.014 |  | 0.846 |  |
| Residual | 5.303 |  | 0.058 |  | 2.459 |  |

<sup>a</sup> Annual dose of N fertilization as numeric variable: 0, 6.25, 12.5, and 25 kg N ha<sup>-1</sup> a<sup>-1</sup>
<sup>b</sup> Temperature as categorial variable. Baseline: 22°C
<sup>c</sup> August vs. June
<sup>d</sup> Days of storage in water before measurement

*A*_max_ was significantly reduced by increases in temperature (Tables 4, S1; Fig. 4a). Furthermore, *R*_d_ was steadily intensified with increasing temperature (Table S1; Fig. 4b). LCP increased to higher values of PPFD with increasing temperature; while this effect was highly significant (*P*≤0.001), high residual variance in the mixed-effects model and the very low *R*^2^ of 0.07 indicated that LCP mostly depended on other factors than temperature, N fertilization, and month of sampling (Table S1). LCP was relatively stable at 25 and 30°C, but increased sharply from 35 to 40°C (Fig. 4c).

**Fig. 4.**
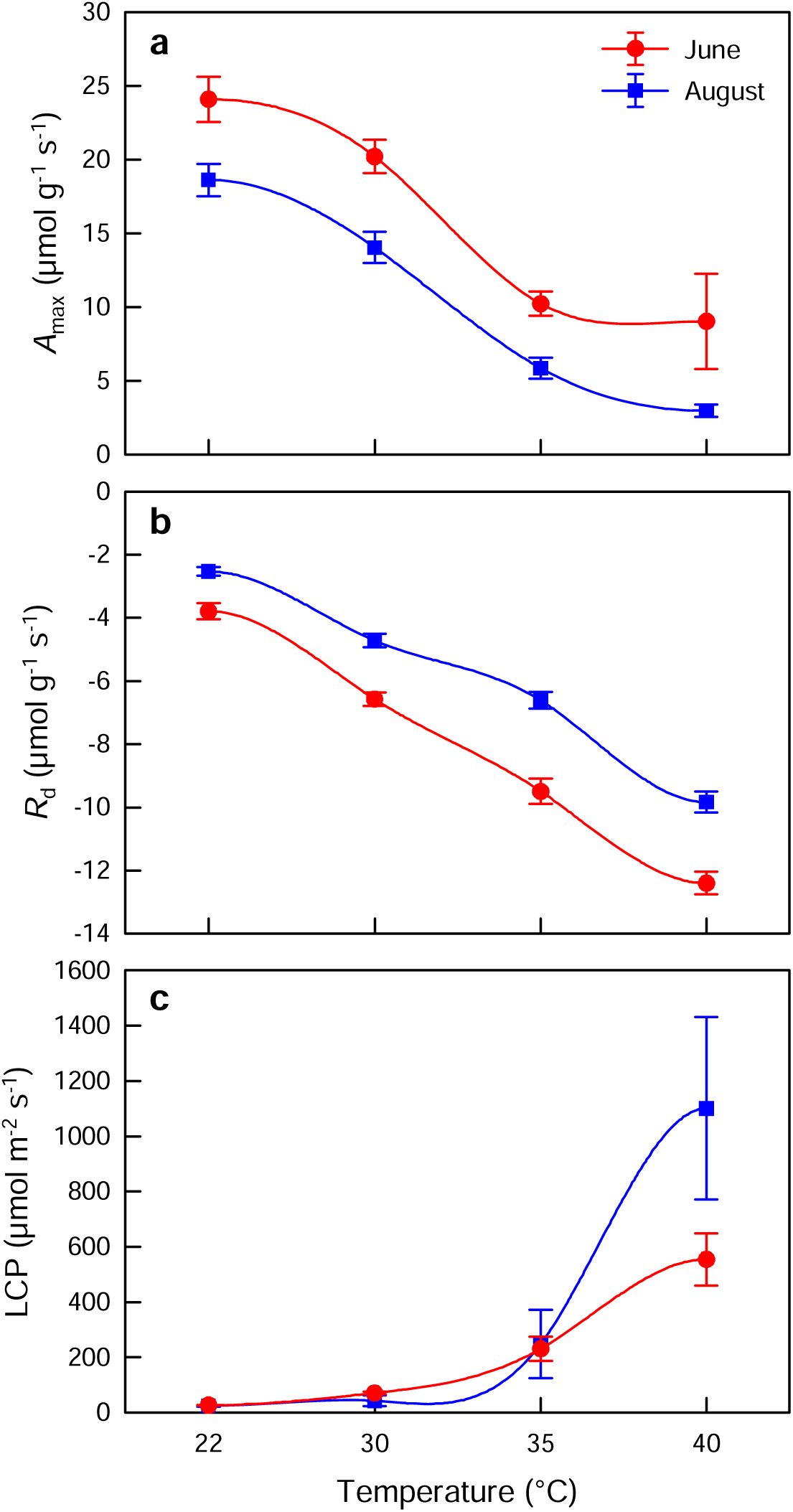
(a) Photosynthetic capacity (*A*_max_), (b) dark respiration (*R*_d_), and (c) light compensation point (LCP) in *Larix sibirica* at different temperatures from 22–40°C in June and August. N fertilization had no significant effect on these parameters.

### 3.3. Early vs. late summer

The month of sampling had significant effects on almost all parameters. It caused opposite effects on *g*_s_ and *E* in the mixed-effects model (Table 2). In June, *g*_s_ was significantly lower, but *E* was higher than in August. *A* and *A*_max_ were both lower in August than in June (Tables 3, 4; Fig. 3). The effect of August vs. June on WUE_i_ and WUE_inst_ was positive (Table 3). *R*_d_ was higher in August than in June (Table 4). LCP was not significantly influenced (Table S1).

## 4. Discussion

### 4.1. Fertilization effects on stomatal conductance

The results substantiate that tree water relations were influenced by N availability in the studied drought-limited southern boreal forests. Consistent with our first hypothesis, both stomatal conductance (*g*_s_) and transpiration (*E*) increased in response to the fertilization of *L. sibirica* forests with 6.25–25 kg N ha^−1^ a^−1^. This result obtained in the third year of N fertilization in the Dry Boreal Forest Long-Term Nitrogen Fertilization Experiment (DryBoreal-LTNE) provides direct experimental evidence that confirms earlier results where an effect of N availability on *g*_s_ of *L. sibirica* was postulated based on stable isotope analyses (δ^13^C, δ^15^N) along spatial gradients of N supply due to local differences in forest grazing intensity (Dulamsuren & Hauck 2021).

Mechanisms how N availability controls *g*_s_ are known in great detail on even a molecular level (Guo et al. 2003; Tsay et al. 2007; Fang et al. 2021). Increases in *g*_s_ with increasing N supply have also early been demonstrated in herbaceous plants and cotton seedlings (Radin & Ackerson 1981; Ghasghaie & Saugier 1989; Broadley et al. 2001) and more recently in potted tree saplings (Zhu et al. 2020; F. Feng et al. 2026b). In our study, we could prove this relationship in drought-limited mature boreal forests from a region devoid of any significant atmospheric N deposition from anthropogenic sources (Rubin et al. 2023; Mgelwa et al. 2024). Our results from highly continental boreal climate match with increased values of canopy conductance (*G*_s_) due to N fertilization that were recently published from a long-term experiment in a boreal *Pinus sylvestris* forest of northern Sweden under a much moister boreal climate (L. Chen et al. 2026). In a study from a southern boreal *L. gmelinii* forest of northern China, Liu et al. (2022) found no significant effect of N fertilization on *g*_s_, which is probably attributable to the application of a much higher annual N dose of 75 kg N ha^−1^ a^−1^. Such failure to generate statistically significant positive responses of *g*_s_ due to N fertilization was also observed in experiments with temperate conifers in the field (Domec et al. 2009) or in pot experiments at high N doses (Green & Mitchell 1992; Paligi et al. 2024) of up to 60 or even 112 kg N ha^−1^ a^−1^.

### 4.2. Utilization of increased g_s_ for increasing the photosynthetic C gain

Under the experimental conditions of our gas exchange measurements, the increases in *g*_s_ could be translated into increased net photosynthesis (*A*) in agreement with our second hypothesis. In the lowest fertilization treatment N1 with an annual N dose of 6.25 kg N ha^−1^ a^−1^, however, any potential increases in gross photosynthesis were masked by the strong increase in dark respiration (*R*_d_). This pattern can be explained with the high metabolic costs of N acquisition at low concentrations of plant-available N (Garcia et al. 2023). Nevertheless, slight, but significant improvements in water-use efficiency (WUE) already at 6.25 kg N ha^−1^ a^−1^ indicated that even gross photosynthesis benefitted from the increased N availability in treatment N1.

The annual N dose of 12.5 kg N ha^−1^ a^−1^ (treatment N2) clearly represented the optimum N supply for the photosynthetic C gain in our experiment. In N2, we observed the highest values of *A*, WUE_i_, and WUE_inst_. However, we also found the highest photosynthetic capacity (*A*_max_) in N2, which thus only partly confirmed our third hypothesis. This increase in *A*_max_ demonstrates that the increased CO_2_ assimilation was not only the result of increases in the intercellular CO_2_ concentrations in the needles due to N-induced increases in *g*_s_, but that the additional N was used to build up higher capacities for CO_2_ assimilation in the dark reaction (e.g. by increasing the amount of Rubisco) and perhaps also by synthesizing addition chlorophyll for the light reaction (Lu et al. 2022). Since we did not measure chlorophyll concentrations, assuming the latter response remains speculative.

That *A* decreased towards the high N dose from treatment N2 to N3 is often a sign of increased respiration (Vose & Ryan 2002; Bosc et al. 2003). Our data are not conclusive to provide a detailed explanation. Yet this decline is not necessarily the sign of some detrimental effect, for instance, for the respiratory effort for NH_4_^+^ detoxification (Bittsánszky et al. 2015), but can also be the consequence of increased respiration for growth and increased production of non-structural carbohydrates (NSC) (Amthor 2000; Collalti et al. 2020; Oswald & Aubrey 2024). Not all of this respiration might be visible as an increase in *R*_d_, and moreover, we did unfortunately not find a powerful model for *R*_d_. Therefore, we cannot draw a final conclusion from our data whether the decline of *A* at 25 kg N ha^−1^ a^−1^ was already a sign of overfertilization or indicated an increase in growth and maintenance respiration (Amthor 2000; Uscola et al. 2015).

### 4.3. Effects of temperature and changes during the growing season

The decreases of *A* and WUE with increasing temperature from 22–40°C reflect the increasing deactivation of Rubisco at high temperatures (Salvucci & Crafts-Brandtner 2004; Sharkey 2005) and the exponential increase of respiration in this temperature range (Scafaro et al. 2021). These declines match with the expectation for a boreal forest tree of a thermal optimum of *A* that is clearly below 30°C (Vygodskaya et al. 1997; Dusenge et al. 2023). The reduction of *g*_s_ at high temperatures is most likely due to an increase in VPD, as relative humidity was kept constant during the measurements (Merilo et al. 2018; Xu et al. 2025; Yang et al. 2026). The slightly lower reduction of *g*_s_ at 40°C than at 35°C, combined with the sharp increase in *E* at 40°C, can be interpreted as the onset of thermal decoupling of *g*_s_ from *A* around 40°C, which is an emergency measure of transpirative cooling to avoid heat damage regardless of the water supply (Marchin et al. 2023; Wang et al. 2026).

Declines in *A* during the growing season can be explained by reductions in *A*_max_ due to the accumulation of damage as the result of photo-oxidative stress (Stock et al. 2026). The significant loss of *A*_max_ within just 2 months is most likely the consequence of the frequent drought stress in the highly continental southern boreal forests of Mongolia. Drought is well-known to increase photo-oxidative stress (Werner et al. 2002; Wujeska et al. 2013; Zahra et al. 2023) and can thus be assumed to further reduce the annual C gain in the short growing season of Mongolia’s boreal forests by fostering photodestruction in addition to the immediate reductions in CO_2_ assimilation during drought due to stomatal closure (Dulamsuren et al. 2009a, 2023b).

### 4.4. Role of eCO_2_-induced oligotrophication and land use for Mongolia’s boreal forests

Combined with results of Dulamsuren & Hauck (2021) – also derived from drought-limited *L. sibirica* forests of Mongolia – the present findings suggest that N availability play a significant role for C and tree water relations not only in cold-limited boreal forests, but also in drought-limited systems. This insight is important, as more and more boreal forests switch from temperature limitation to drought limitation due to climate change (Babst et al. 2019; Mirabel et al. 2023; Sang et al. 2026). In our CO_2_/H_2_O gas exchange measurements, we analyzed the effects of long-term N fertilization in combination with short-term responses to climatic conditions, which were varied during the measurements in order to obtain direct evidence for the influence of N supply on *g*_s_ and *A*. Using combined dendrochronological and stable isotope analyses, Dulamsuren & Hauck (2021) could show an even stronger influence of N availability on the variability of radial stem increment in *L. sibirica* than of drought. The tight relationship to N availability might explain the often low *R*^2^ values in dendrochronological climate-response analyses, because these analyses do only rarely include N availability or N deposition as an additional factor (Hauck et al. 2012; Z. Chen et al. 2026).

Against the background of an eCO_2_-induced global oligotrophication of remote boreal forests that is assumed by an increasing number of ecologists (Craine et al. 2018; Mason et al. 2022a; Bassett et al. 2026) – though not by all authors (Hiltbrunner et al. 2019; Olff et al. 2022) – the long-term responses of boreal forest ecosystems to global change have to be regarded from a broader perspective than usually done: In addition to effects of drought (Dulamsuren et al. 2010; Babst et al. 2019) and heat (Rao et al. 2023; Dulamsuren et al. 2026) more emphasis should be placed on the study of interactions of drought stress and N given the expectation that the availability of N will most likely further decrease in boreal forests (Bassett et al. 2026) concomitant to increased drought stress (Zhang et al. 2020; Mirabel et al. 2023). So far, most work on the interaction of N availability and tree water relations is from temperate forests (Audisio et al. 2024; Dietrich et al. 2024; Hauck & Dietrich 2025; Tresch et al. 2025) and from moist boreal forests (Palvi et al. 2025; L. Chen et al. 2026), but much less in known from boreal forest ecosystems in regions with strongly continental climate under drought, which might represent larger boreal forest areas under future climate (Mirabel et al. 2023; Runge et al. 2025).

In the special case of Mongolia’s boreal forests, forest grazing counteracts oligotrophication and generates spatial variability in N availability (Dulamsuren & Hauck 2021). This way, Mongolia’s pastoral nomadism, the socioecological impact of which on forests has previously been only discussed in terms of livestock feeding damage in the forest regeneration (Khishigjargal et al. 2013; Juřička et al. 2020) and of fuel and construction wood consumption by the herders (Hauck et al. 2024), attains a role in modifying the tolerance of Mongolia’s southern boreal forests to climate change, where livestock dung counteracts eCO_2_-induced reductions in plant-available N and thus increases *g*_s_ due to N-induced stomatal opening in *L. sibirica*. This might be beneficial in periods of moderate drought due to increased *A* (this study) and growth (Dulamsuren & Hauck 2021), but might be harmful if drought intensifies due to high tissue water loss.

## 5. Conclusions

Our CO_2_/H_2_O measurements demonstrated that increased N availability (fertilization with 6.25–25 kg N ha^−1^ a^−1^ for 3 years) interfered with the stomatal control by soil moisture and VPD in *L. sibirica*, which dominates the drought-limited southern boreal forests in Mongolia. Increased N availability increased stomatal conductance (*g*_s_), which was utilized for increases in net photosynthesis (*A*) and has thus the potential to retard stomatal closure and increase forest productivity during moderate droughts. However, if drought intensifies, it is plausible to assume that N-induced increases in *g*_s_ increase the risk of tree mortality, as the relative tissue water content will decrease more strongly than at lower N levels. This interaction of nitrogen and tree water relations is significant, because remote boreal forests are assumed to have been depleted in plant-available N since centuries due to elevated atmospheric CO_2_ (eCO_2_), on the one hand (Bassett et al. 2026). On the other hand, forest grazing has been shown to increase N availability in Mongolia, thus locally counteracting the oligotrophication of Mongolia’s boreal forests. The drought vulnerability of these southern boreal forests does thus not only depend on climate and soil conditions, but in addition on the land use practices by pastoral livestock herders and on the global rise of the atmospheric CO_2_ concentration due to fossil fuel combustion. Direct heat effects additionally influence the CO_2_ assimilation, as *A* was reduced at temperatures ≥30°C. Our study shows that focusing on tree hydraulics and tree water status in response to meteorological variables alone does only explain part of the response of these dry southern boreal forests to global change.

## Acknowledgments

We are thankful to the Forest Office (Mr. D. Batbaatar, Mr. B. Tseveldorj) in Jargalant, Arkhangai district, Mongolia for multiple support during field work and the permission to set up the DryBoreal Forest-LTNE field sites.

## Funding information

German Research Foundation (Deutsche Forschungsgemeinschaft). Grant/Award Number: 539975714 approved to Choimaa Dulamsuren (Du 1145/5-1). Academic Society of Freiburg im Breisgau (Wissenschaftliche Gesellschaft). Travel grant approved to Choimaa Dulamsuren.

## CRediT authorship contribution statement

**Choimaa Dulamsuren:** Conceptualization, Methodology, Investigation, Data curation, Formal analysis, Writing – original draft, Writing – Review & Editing, Funding acquisition. **Dugersuren Amarjargal:** Investigation. **Dashtogtokh Bazar:** Investigation. **Tumendelger Uitumen:** Investigation. **Tungalag Munkhzul:** Investigation. **Davaadorj Saindovdon:** Investigation. **Batbold Byambatsogt:** Investigation. **Erdenechuluun Naranbayar:** Investigation. **Ganbaatar Batsaikhan:** Investigation, **Markus Hauck:** Conceptualization, Investigation, Writing – original draft, Writing – Review & Editing.

## Conflict of interest

The authors declare no conflict of interest.

## Data availability statement

Data will be made available on Zenodo after acceptance.

---

**Table S1.** Linear mixed-effects model analyzing effects of N fertilization, temperature, and month on photosynthetic capacity (*A*_max_), dark respiration (*R*_d_), and light compensation point (LCP) in *Larix sibirica*.

|  | *A*<sub>max</sub> (µmol g<sup>-1</sup> s<sup>-1</sup>) |  | *R*<sub>d</sub> (µmol g<sup>-1</sup> s<sup>-1</sup>) |  | LCP (µmol m<sup>-2</sup> s<sup>-1</sup>) |  |
|---|---|---|---|---|---|---|
|  | Estimate±SE | *P* | Estimate±SE | *P* | Estimate±SE | *P* |
| Fixed effects: |  |  |  |  |  |  |
| (Intercept) | 15.885±0.843 | **<0.001** | -8.104±0.318 | **<0.001** | 221.25±70.29 | **0.008** |
| Nitrogen<sup>a</sup> | 0.631±0.535 | 0.24 | -0.001±0.102 | 0.99 | 33.11±46.87 | 0.48 |
| Temperature | -6.124±0.532 | **<0.001** | -2.908±0.100 | **<0.001** | 272.95±46.87 | **<0.001** |
| Month<sup>b</sup> | -5.653±1.080 | **<0.001** | 2.236±0.214 | **<0.001** | 134.51±93.65 | 0.15 |
| Nitrogen × Temperature | -0.291±0.532 | 0.58 | 0.040±0.100 | 0.69 | 34.85±46.86 | 0.46 |
| Conditional *R*<sup>2</sup> | 0.26 |  | 0.68 |  | 0.07 |  |
| Marginal *R*<sup>2</sup> | 0.25 |  | 0.65 |  | 0.07 |  |
| Random effects: |  |  |  |  |  |  |
| Site | 0.402 |  | 0.380 |  | 53.13 |  |
| Storage<sup>c</sup> | 0.778 |  | 0.556 |  | 0.00 |  |
| Residual | 11.610 |  | 2.181 |  | 1024.78 |  |

<sup>a</sup> Annual dose of N fertilization as numeric variable: 0, 6.25, 12.5, and 25 kg N ha<sup>-1</sup> a<sup>-1</sup>
<sup>b</sup> August vs. June
<sup>c</sup> Days of storage in water before measurement

**Table S2.**
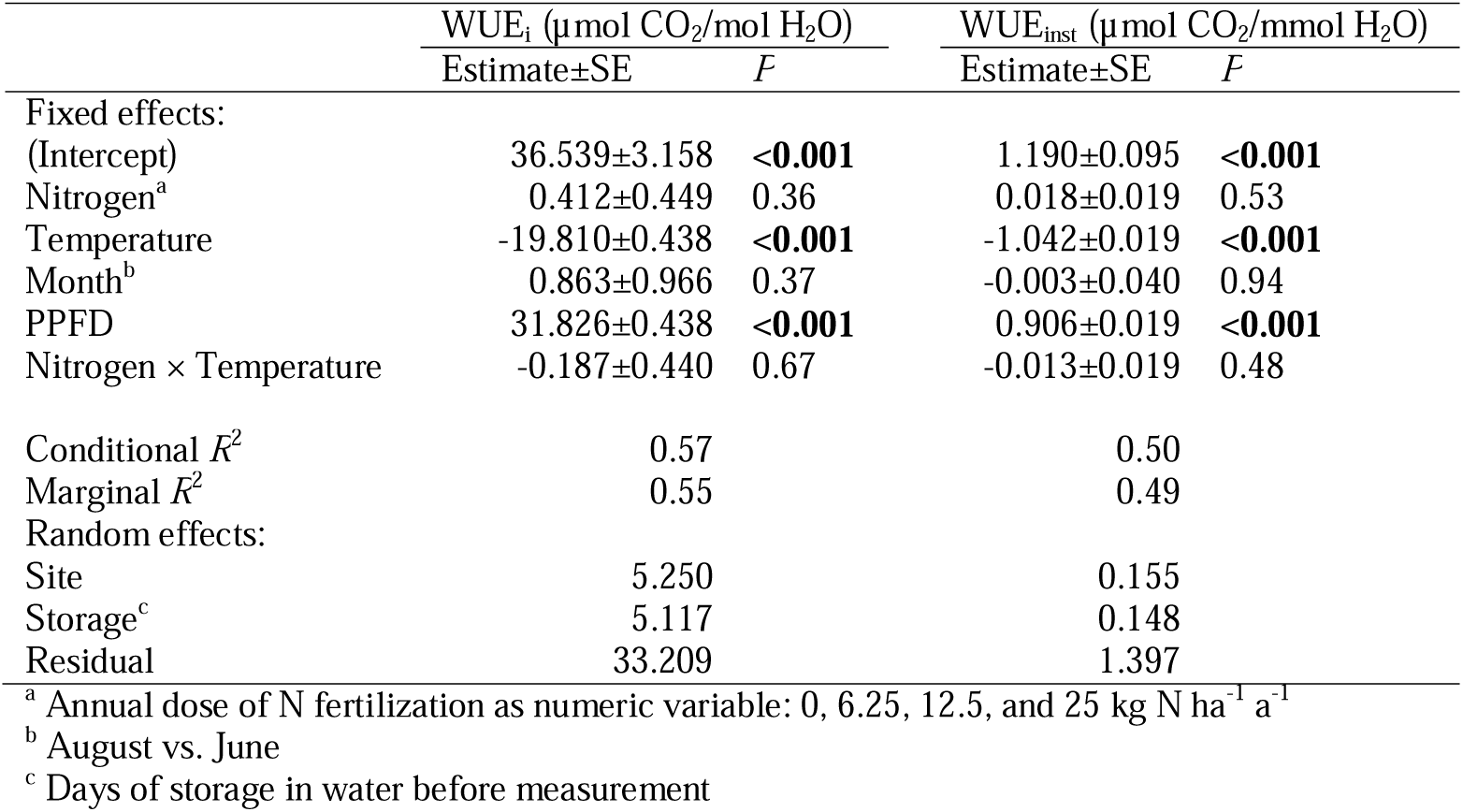
Linear mixed-effects model analyzing effects of N fertilization, temperature, and month on intrinsic (WUE_i_) and instantaneous (WUE_inst_) water-use efficiency in *Larix sibirica*^a^.

|  | WUE<sub>i</sub> (µmol CO<sub>2</sub>/mol H<sub>2</sub>O) |  | WUE<sub>inst</sub> (µmol CO<sub>2</sub>/mmol H<sub>2</sub>O) |  |
|---|---|---|---|---|
|  | Estimate±SE | *F* | Estimate±SE | *F* |
| Fixed effects: |  |  |  |  |
| (Intercept) | 36.539±3.158 | <**0.001** | 1.190±0.095 | <**0.001** |
| Nitrogen<sup>a</sup> | 0.412±0.449 | 0.36 | 0.018±0.019 | 0.53 |
| Temperature | -19.810±0.438 | <**0.001** | -1.042±0.019 | <**0.001** |
| Month<sup>b</sup> | 0.863±0.966 | 0.37 | -0.003±0.040 | 0.94 |
| PPFD | 31.826±0.438 | <**0.001** | 0.906±0.019 | <**0.001** |
| Nitrogen × Temperature | -0.187±0.440 | 0.67 | -0.013±0.019 | 0.48 |
| Conditional *R*<sup>2</sup> | 0.57 |  | 0.50 |  |
| Marginal *R*<sup>2</sup> | 0.55 |  | 0.49 |  |
| Random effects: |  |  |  |  |
| Site | 5.250 |  | 0.155 |  |
| Storage<sup>c</sup> | 5.117 |  | 0.148 |  |
| Residual | 33.209 |  | 1.397 |  |

<sup>a</sup> Annual dose of N fertilization as numeric variable: 0, 6.25, 12.5, and 25 kg N ha<sup>-1</sup> a<sup>-1</sup>
<sup>b</sup> August vs. June
<sup>c</sup> Days of storage in water before measurement

## Notes

### Competing Interest Statement

The authors have declared no competing interest.

